# Garcinia cambogia extract removes calcium oxalate kidney stones in both genetic and non-genetic *Drosophila* models of nephrolithiasis

**DOI:** 10.1101/477570

**Authors:** Qiuxia Fan, Xiaoming Feng, Xizhen Hong, Siqiao Gong, Jianwei Tian, Fanfan Hou, Fujian Zhang

## Abstract

Kidney stone formers with family history have a high rate of stone recurrence after kidney stone removal surgery and there is no effective medication available for treatment. Here, we show that *Garcinia cambogia* extract (GCE) efficiently removes calcium oxalate kidney stones from Malpighian tubules in both genetic and non-genetic *Drosophila* models of nephrolithiasis, and hydroxycitrate -a major component of GCE, directly dissolves calcium oxalate stones in *Drosophila* Malpighian tubules *ex vivo*. Our study discovers a potential novel therapeutic strategy for the clinical treatment of nephrolithiasis and suggests that clinical-grade *Garcinia cambogia* extract could be used to treat patients with nephrolithiasis in the future.

## Introduction

The incidence of nephrolithiasis, also known as kidney stone, is increasing globally and puts a huge burden on health care system gradually. Kidney stones often cause hematuria and severe pain in the groin, abdomen, or flank. Calcium-containing kidney stones are the most common type of kidney stone and account for more than 90% of renal stone diseases (Gee *et al.*, 2016). Although almost 50% of calcium oxalate (CaOx) stone formers are idiopathic(Ramaswamy *et al.*, 2015, Chi and Stoller, 2011), kidney stones have long been thought to be the result of the interaction between genetic and environmental factors, including dehydration stemming from low fluid intake and high dietary intake of animal protein and salts. Meanwhile, more than 30 genes up to date have been identified as novel monogenic causes of kidney stone disease using whole exome sequencing (Gee et al., 2016, Halbritter *et al.*, 2015, Braun *et al.*, 2016, Daga *et al.*, 2018, Sayer, 2017, Dowen *et al.*, 2014). Effective medications are generally lacking, and the most advanced treatment for kidney stones is minimally invasive kidney stone surgical procedures, such as shockwave lithotripsy (SWL), ureteroscopy and laser lithotripsy and percutaneous nephrolithotomy (PCNL) depending on the size and type of stones. Despite great improvement in surgical techniques used to remove kidney stones in the last two decades, patients very often experience a high rate of stone recurrence after surgery, especially those who are inherited cases. The high rate of stone recurrence and the surgical burden associated with these patients demand the discovery of new drugs that could dissolve kidney stones *in situ*. However, no major progress of new drug discovery has been made to remove or prevent kidney stone formation in the last 30 years, mainly due to the lack of ideal animal models feasible for high-throughput drug screening.

The *Drosophila* excretory system is composed of nephrocytes and Malpighian tubules(Zhang and Chen, 2014). Our previous study showed that the *Drosophila* nephrocyte shares remarkable similarity with the glomerular podocyte for protein ultrafiltration, and the renal proximal tubule for protein reabsorption(Gee *et al.*, 2015, Zhang and Chen, 2014, Zhang *et al.*, 2013, Zhang *et al.*, 2013). On the other hand, the *Drosophila* Malpighian tubule shares striking similar features with mammalian renal tubules and collecting ducts in terms of cell composition, anatomical structure and physiological function(Dow and Romero, 2010). There are two types of cells in *Drosophila* Malpighian tubule, the principal cells and stellate cells which contain many ion and organic solute transporters (Miller *et al.*, 2013). The principal cells are the major tubular cell type (∼80%) through which cations and organic solutes are transported. The stellate cells are the minor tubular cell type (∼20%) through which chloride ion and water flow, and interspersed at regular intervals with the principal cells. They generate urine through active transport of ions, water and organic solutes from the hemolymph into the Malpighian tubule lumen(Dow and Romero, 2010). It has been shown that *Drosophila* Malpighian tubule is a powerful translational model system to study the pathogenesis of human nephrolithiasis, because one can easily generate the disease model, observe stones and perform genetic manipulation (Chen *et al.*, 2017, Wu *et al.*, 2014, Miller et al., 2013, Hirata *et al.*, 2012a, Knauf and Preisig, 2011, Chi *et al.*, 2015). Calcium oxalate stones are formed in Malpighian tubule with the addition of lithogenic agents in fly food within two weeks and can be directly examined under polarized light microscopy (Chen *et al.*, 2011). Slc26a6 functions as a Cl^-^/HCO3^-^ exchanger and regulates oxalate secretion in human renal tube. Mutations of Slc26a6 have been identified from patients with nephrolithiasis. It also has been shown that RNAi knockdown of dPrestin specifically in principal cells, the *Drosophila* homolog of Slc26a6, led to decreased calcium oxalate stone formation in Malpighian tubules(Landry *et al.*, 2016). These results strongly suggest that fruit fly could be an ideal genetic kidney stone disease model to screen novel genes involved in the pathogenesis of nephrolithiasis and validate the function of candidate genes identified from patients with nephrolithiasis *in vivo*(Hirata et al., 2012a).

The *Drosophila* calcium oxalate stone model has been used to screen traditional Chinese medicinal plants for the treatment of nephrolithiasis (Ho *et al.*, 2013). Chen *et al* showed that commercial juices, such as apple, cranberry, orange, and pomegranate juices, failed to prevent calcium oxalate crystal formation in *Drosophila* nephrolithiasis model. Wu *et al* showed that some traditional Chinese Medicine (TCM) herbs have a potential antilithic effect, but it is still unclear whether they can directly dissolve kidney stones *in situ* to prevent kidney stone recurrence. In the last 30 years, no major progresses have been made in this field. *Garcinia cambogia* is a tropical fruit that grows in Southeast Asia and has been historically used for cooking in India. *Garcinia cambogia* extract containing 60% hydroxycitric acid (HCA), is a popular weight-loss supplement and sold at most health supplement and drug stores. Chung *et al* showed that HCA induces dissolution of the calcium oxalate crystal *in vitro*, suggesting that *Garcinia cambogia* has the potential as a novel treatment for calcium oxalate kidney stone(Chung *et al.*, 2016). In this study, we used the *Drosophila* genetic nephrolithiasis model to test whether *Garcinia cambogia* and hydroxycitrate can prevent the formation of calcium oxalate stones *in vivo*. We found that *Garcinia cambogia* extract prevented calcium oxalate kidney stone formation and completely removed calcium oxalate kidney stones preformed in adult renal tubules *in vivo*. To further elucidate the molecular mechanism through which GCE removed calcium oxalate stones from renal tubules, we showed that hydroxycitrate - the major component of GCE - directly dissolved calcium oxalate stones in renal tubules *ex vivo*. Our data strongly suggest that clinical-grade *Garcinia cambogia* extract could be used to remove calcium oxalate renal stones in patients with nephrolithiasis.

## RESULTS

### *Garcinia cambogia* extract efficiently prevents calcium oxalate kidney stone formation in *Drosophila* renal tubules *in vivo*

To test whether *Garcinia cambogia* can prevent the formation of calcium oxalate stones *in vivo*, w^1118^ wild type flies were reared in fly food containing 0.1% NaOx and different concentrations of GCE for one week, and then the effect of GCE on the formation of kidney stone in renal tubules was examined. We found that *Garcinia cambogia* extract prevented calcium oxalate kidney stone formation in adult renal tubules *in vivo* in a concentration-dependent manner (Fig.1A-D). Compared with hydroxycitrate or citrate, *Garcinia cambogia* extract prevented the formation of calcium oxalate stone in *Drosophila* renal tubules at a very low concentration of 0.3% (Fig.1E). On the other hand, HCA or citric acid (CA) only partially blocked the formation of calcium oxalate stone in *Drosophila* renal tubules at a high concentration of 1.5% (Fig.1E), and both of them prevented calcium oxalate kidney stone formation in *Drosophila* renal tubules at a very high concentration of 3% (Fig.1E). These results indicate that *Garcinia cambogia* extract are a better reagent to prevent calcium oxalate stone formation in kidney stone disease models *in vivo* than citric acid which is widely used in clinic.

**Figure1.**
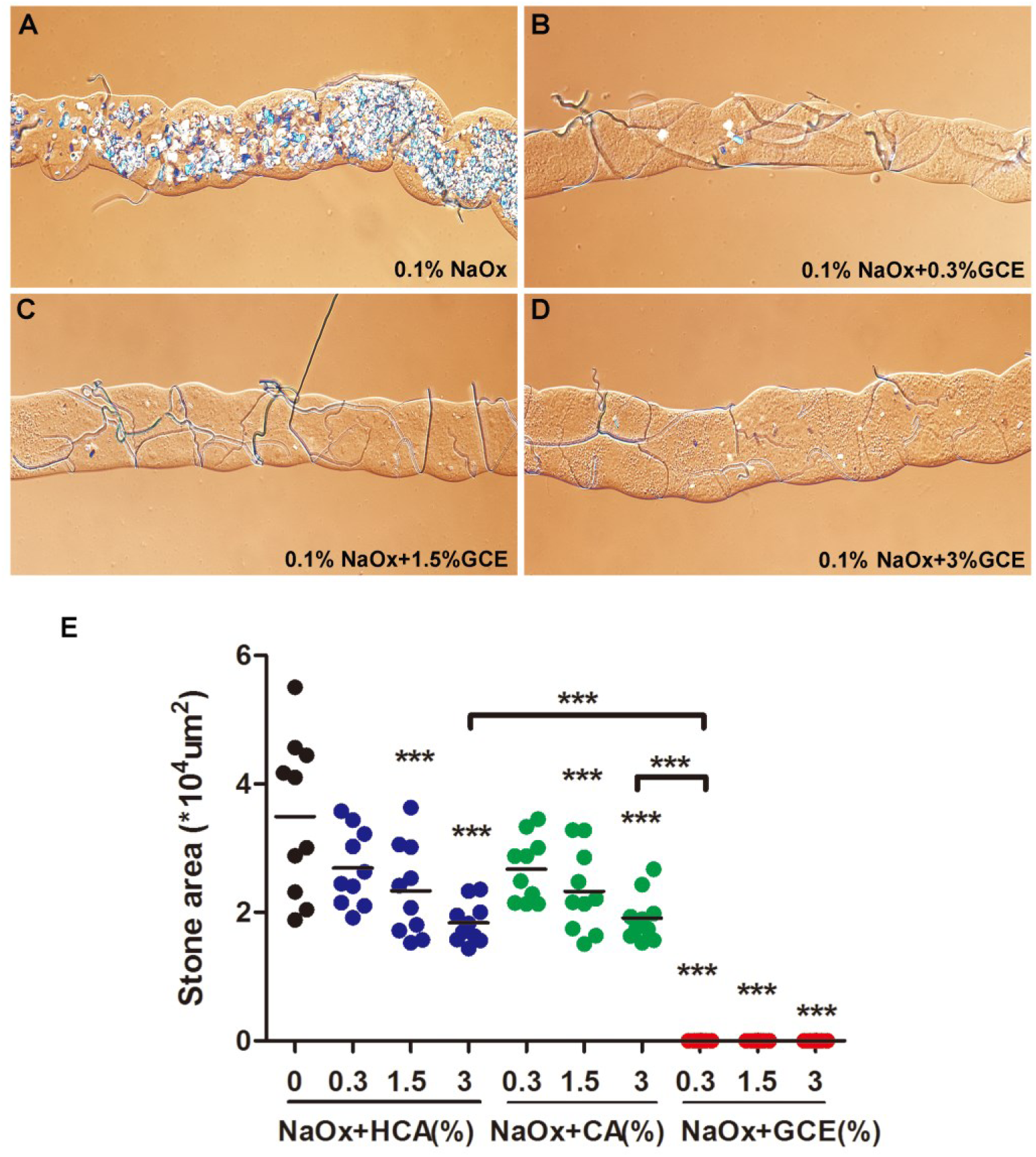
*Garcinia cambogia* extract prevents calcium oxalate kidney stone formation in *Drosophila* renal tubules. Representative images of the effect of GCE on the formation of CaOx stone in adult malpighian tubules (A-D). Calcium oxalate kidney stone formation in wild type flies reared in fly food containing 0.1% NaOx (A), 0.1% NaOx+0.3% GCE (B), 0.1% NaOx+1.5% GCE (C), 0.1% NaOx+3% GCE (D). 10 pairs of Malpighian tubules from 5 flies per genotype were dissected and analyzed. Total area of CaOx stones in Malpighian tubule was measured in the whole field of view (700umx100um, 20 x magnification). The results are expressed as mean ± SD. One-way ANOVA was performed to analyze the data and Bonferroni’s multiple comparison was performed to compare all pairs of columns. Statistical significance was defined as p<0.05 (*p<0.05, **p<0.005, ***p<0.0005). (E) Comparison of the effect of GCE, HCA and CA on calcium oxalate kidney stone formation. GCE totally prevented calcium oxalate kidney stone formation in adult renal tubules at 0.3%,1.5% and 3% compared to control (***p<0.0001); whereas, 3% of HCA or CA only partially prevented calcium oxalate kidney stone formation compared to control (***p<0.0001). 0.3% of GCE is more efficient to prevent calcium oxalate kidney stone formation compared to 3% of HCA or CA (***p<0.0001).

### *Garcinia cambogia* extract completely removes calcium oxalate kidney stones from adult *Drosophila* renal tubules *in vivo*

Previous study showed that hydroxycitric acid (HCA) can induce the dissolution of the calcium oxalate crystal *in vitro.* We reasoned that *Garcinia cambogia* extract (GCE) which contains 60% HCA could play a similar role *in vivo*. To test the effect of GCE on the removal of calcium oxalate renal stones *in vivo*, we first developed a calcium oxalate kidney stone disease model by feeding wild type flies with fly food containing 0.3% NaOx for one week. Flies were then transferred to fly food containing 0.1%NaOx and different concentrations of GCE for another week, and then the effect of GCE on the removal of kidney stone from renal tubules was examined. As shown in Figure 2, treatment with different concentrations of GCE for one week removed calcium oxalate kidney stones from adult renal tubules in a concentration-dependent manner (Fig.2A-D). The total stone area decreased by about 93% in renal tubules of flies reared in fly food with 0.3% GCE compared to flies reared in 0.1% NaOx only. No stones were left in renal tubules of flies reared in fly food with 1.5% (Fig. 2E) or 3% GCE (Fig. 2E). Compared to GCE, hydroxycitrate removed calcium oxalate kidney stone formation at the concentrations of 0.3%, 1.5% with much less efficiency (Fig.2E). The total stone area decreased by about 50% in renal tubules of flies reared in fly food with 0.3% HCA compared to flies reared in 0.1% NaOx only. Citric acid could only partially remove calcium oxalate stone in *Drosophila* renal tubules at a very high concentration of 3%. The total stone area decreased by 20% compared to the flies in control group (Fig.1E). Our results strongly suggest that *Garcinia cambogia* extract could be used to efficiently remove calcium oxalate renal stones in calcium oxalate nephrolithiasis patients.

**Figure2.**
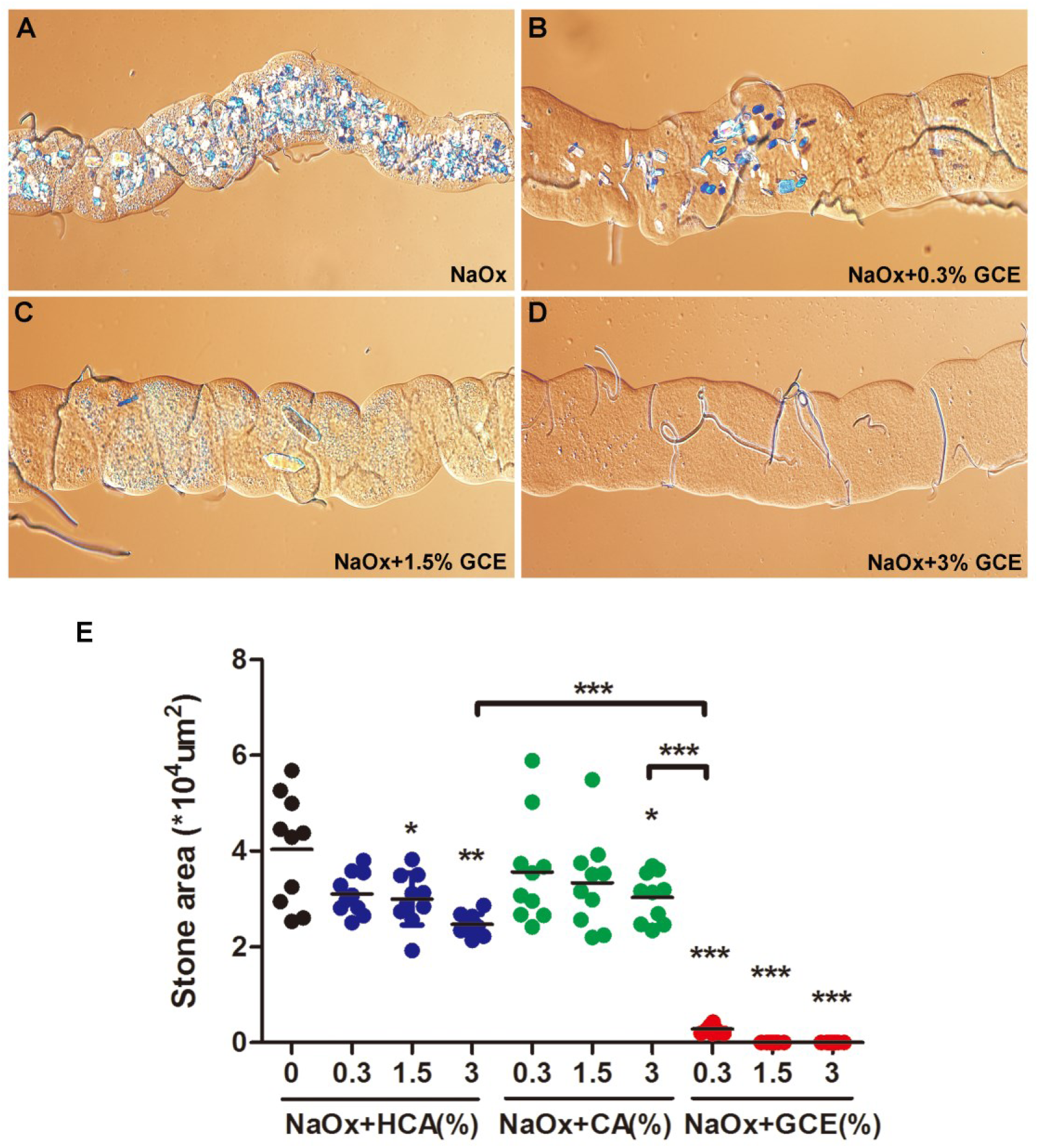
*Garcinia cambogia* extract completely removes calcium oxalate kidney stones from *Drosophila* renal tubules *in vivo*. Representative images of the effect of GCE on the removal of CaOx stone in adult Malpighian tubules (A-D). Wild type flies were reared in fly food containing 0.3% NaOx for one week and then transferred to new fly food containing 0.1% NaOx (A), 0.1% NaOx+0.1% GCE (B), 0.1% NaOx+0.5% GCE (C), 0.1% NaOx+1% GCE (D) for one week. 10 pairs of Malpighian tubules from 5 flies per genotype were dissected and analyzed. Total area of CaOx stones in Malpighian tubule was measured in the whole field of view (700umx100um, 20 x magnification). The results are expressed as mean ± SD. One-way ANOVA was performed to analyze the data and Bonferroni’s multiple comparison was performed to compare all pairs of columns. Statistical significance was defined as p<0.05 (*p<0.05, **p<0.005, ***p<0.0005). (E) Comparison of the effect of GCE, HCA and CA on calcium oxalate kidney stone removal. Almost all calcium oxalate kidney stones in adult renal tubules were removed in the presence of 1.5% or 3% GCE compared to control (***p<0.0001), whereas, only 40% of calcium oxalate kidney stones were removed even in the presence of 3% HCA or CA(**p<0.005 or *p<0.05). 0.3% of GCE is more efficient to remove calcium oxalate kidney stone formation compared to 3% of HCA or CA (***p<0.0001).

### *Garcinia cambogia* extract plays a similar role in genetic calcium oxalate kidney stone *Drosophila* model

To test whether *Garcinia cambogia* extract plays a similar role in genetic nephrolithiasis cases, we first developed a genetic calcium oxalate kidney stone *Drosophila* model based on published protocol (Hirata *et al.*, 2012b). We reasoned that if Malpighian tubule is an ideal genetic model for nephrolithiasis, RNAi knock-down of nephrolithiasis-related genes in Malpighian tubule should lead to the formation of calcium oxalate stones. Mutations in the vacuolar-type H^+^-ATPase (v-ATPase) subunit gene ATP6V1B1 and ATP6V0A4 have been identified in recurrent calcium oxalate kidney stone formers with the highest frequency, indicating that v-ATPase is essential to the formation of calcium oxalate kidney stones (Dhayat *et al.*, 2016). The principal cells are the major Malpighian tubular cell type through which cations are transported. Vha55 and Vha100-2 are fly homologs of ATP6V1B1 and ATP6V0A4, and highly expressed in Malpighian tubules (FlyAtlas). UAS-RNAi/Gal4 system has been widely used to knock-down gene expression in a cell-specific manner in *Drosophila*. To specifically silence Vha55 and Vha100-2 genes in Malpighian tubule principal cells, we used the Uro-GAL4 that drives gene expression specifically in the tubular principal cells (Dow and Romero, 2010, Hirata et al., 2012b), and crossed it to the UAS-RNAi lines containing a dsRNA hairpin directed against Vha55 and Vha100-2. We found that RNAi knock-down of vha55 and vha100-2 significantly led to increased formation of calcium oxalate stone in Malpighian tubules compared to control (Uro-Gal4/+)(Figure 3A,D,G). RNAi knockdown of other v-ATPase subunits also resulted in increased calcium oxalate stone formation (Fig.S1). Control or vha100-2/vha55 RNAi knockdown flies were reared in fly food containing 0.3% NaOx for one week. Flies were then transferred to fly food containing 0.1%NaOx and different concentration of GCE (0%, 0.5% and 1%) for another week, and calcium oxalate stone formation was examined in adult Malpighian tubules. As shown in Figure 3, *Garcinia cambogia* extract (GCE) removed calcium oxalate stones preformed in renal tubules of vha55/vha100-2 RNAi knockdown flies in a concentration-dependent manner (Fig. 3A-J). The total stone area in renal tubules was significantly decreased to 12.68% (0.1% NaOx+0.5% GCE) and 0% (0.1% NaOx+1% GCE) compared to flies reared in 0.1% NaOx only. These results showed that *Garcinia cambogia* extract efficiently removed calcium oxalate stones in genetic calcium oxalate kidney stone *Drosophila* model. Altogether, our results strongly suggest that *Garcinia cambogia* extract could be used to remove calcium oxalate stones in both genetic and non-genetic kidney stone disease models.

**Figure3.**
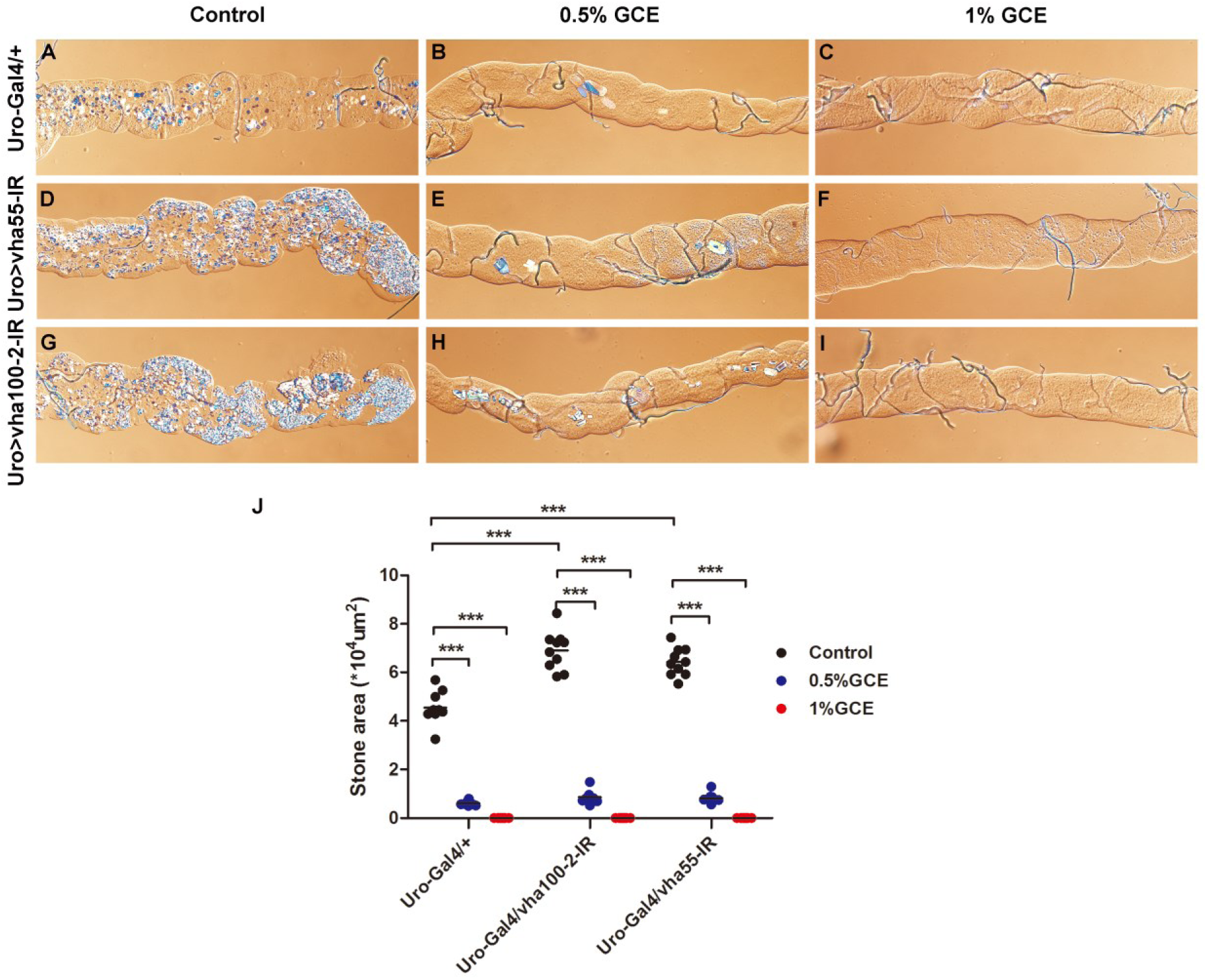
*Garcinia cambogia* extract efficiently removes calcium oxalate kidney stones in genetic nephrolithiasis *Drosophila* model. Representative images of the effect of GCE on the removal of CaOx stone in adult Malpighian tubules of control Uro-Gal4/+ (A-C), Uro-Gal4/vha55 RNAi knockdown (D-F) and Uro-Gal4/vha100-2 RNAi knockdown flies (G-I). Flies reared in fly food containing 0.3% NaOx for one week were transferred to food containing 0.1% NaOx, 0.1% NaOx+0.5% GCE, 0.1% NaOx+1% GCE for one week. 10 pairs of Malpighian tubules from 5 flies per genotype were dissected and analyzed. Total area of CaOx stones in Malpighian tubule was measured in the whole field of view (700umx100um, 20 x magnification). The results are expressed as mean ± SD. Two-way ANOVA grouped analyses was performed to analyze the data and Bonferroni’s post-test was performed to compare replicate means by row. Statistical significance was defined as p<0.05 (*p<0.05, **p<0.005, ***p<0.0005). (J). Comparison of the effect of GCE on the removal of calcium oxalate kidney stones in adult Malpighian tubules of control, vha55 RNAi knockdown and vha100-2 RNAi knockdown flies. RNAi knock-down of vha55 and vha100-2 led to increased formation of calcium oxalate stone in Malpighian tubules compared to control (Uro-Gal4/+)(***p<0.0001). All calcium oxalate kidney stones in adult renal tubules were removed in the presence of 1% GCE in all three groups.

### Hydroxycitrate efficiently dissolves calcium oxalate stones in *Drosophila* renal tubules *ex vivo*

We have showed earlier that GCE can remove calcium oxalate stones from renal tubules *in vivo*. There are two possible pathways for this process: GCE can either facilitate the excretion of stones out of the Malpighian tubule or directly dissolve calcium oxalate stones *in situ*. To further elucidate the molecular mechanism through which GCE removes calcium oxalate stones from renal tubules, we monitored the dissolution of calcium oxalate stones pre-formed in the Malpighian tubule using live-imaging technique. Renal stone dissolution rate was calculated by dividing the remaining stone area by the total stone area before treatment. As shown in Figure 4, HCA directly dissolved calcium oxalate stones in renal tubules *ex vivo* in a concentration- and time-dependent manner. No stones were excreted from the Malpighian tubule during our observation. Fifty percent of CaOx stones were dissolved in 0.5% HCA solution in 2 hours, and were completely dissolved in 1% HCA and 1% CA within 2 hours depending on the original size of stones. There are no difference between HＣＡAND CA. At the same time, we also monitored the effect of *Garcinia cambogia* extract on the dissolution of calcium oxalate stones pre-formed in the Malpighian tubule for 3 hours. We did not observe any impact of GCE on the removal of calcium oxalate stones *ex vivo*, suggesting that the release of free HCA from GCE is essential for its function to dissolve the calcium oxalate stones *in vivo*. GCE added in fly food could be absorbed in midgut and released into the hemolymph as free HCA form, and then secreted into Malpighian tubule lumen where HCA could dissolve calcium oxalate renal stones．Our results strongly support the idea that *Garcinia cambogia* extract removes calcium oxalate renal stones via the direct dissolution of stones by HCA in the renal tubules.

**Figure4.**
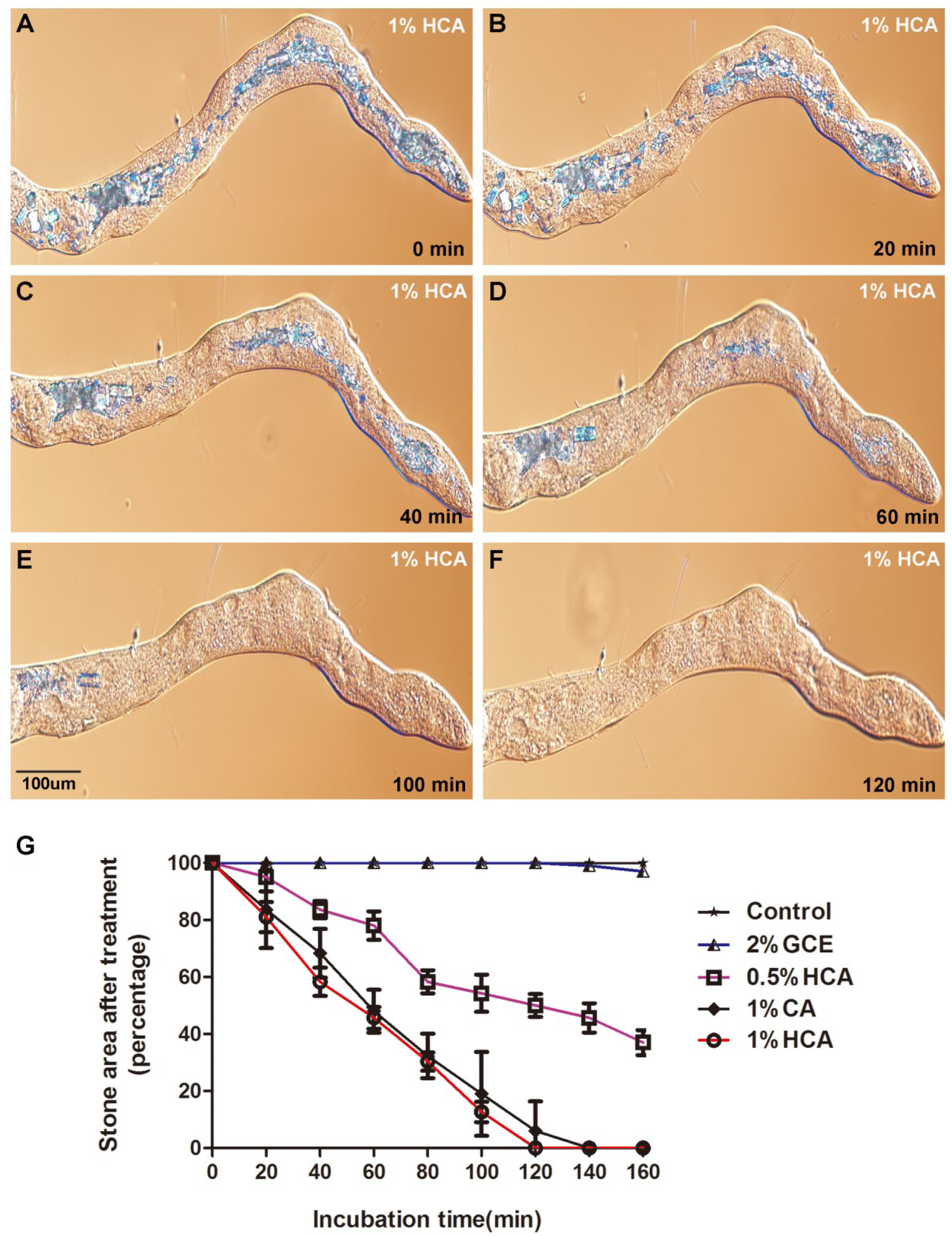
Hydroxycitrate efficiently dissolves calcium oxalate stones in Drosophila renal tubules *ex vivo*. Live images of renal stone dissolution in intact Malpighian tubules treated with HCA solutions ex vivo at different time points (A-F). Wild type flies were reared in fly food containing 0.3% NaOx for one week. Intact Malpighian tubules were dissected from wild type flies reared in fly food containing 0.3% NaOx and treated with HCA, CA or GCE solutions, and renal stone dissolution was monitored using live-imaging. Images were taken at 0min (A), 20min (B), 40min (C), 60min (D), 100min (E), 120min (F). The total area of renal stones was measured using cellSens software. Renal stone dissolution rate was calculated by dividing the remaining stone area by the total stone area at the beginning. The results are expressed as mean ± SD (n=3). (G). HCA and CA efficiently dissolved calcium oxalate stones within 120 minutes. Similar to control group (H2O), GCE had no effect on dissolving calcium oxalate stones. Same scale bars were used in A-F (100um).

## DISCUSSION

Over the past 30 years, no major progress has been made in novel drug discovery for kidney stone treatment because of the lack of ideal animal models feasible for high-throughput drug screening. In this study, we established a genetic nephrolithiasis *Drosophila* model which can be used to screen novel drugs for calcium oxalate renal stones. RNAi knock-down of 5 subunits of v-ATPase using UAS-RNAi/Uro-Gal4 system dramatically enhanced calcium oxalate stone formation in *Drosophila* Malpighian tubules, suggesting that v-ATPase complex in principal cells is essential for calcium oxalate stone formation and *Drosophila* is an ideal model to study the molecular function of genes identified from patients with genetic nephrolithiasis.

*Garcinia cambogia* is a tropical fruit that grows in Southeast Asia and its rind has been widely used for cooking in India. *Garcinia cambogia* extract containing 60% hydroxycitrate is a popular weight-loss supplement sold at most health supplement and drug stores. Chung *et al* has showed that HCA induces dissolution of the calcium oxalate crystal *in vitro*(Chung et al., 2016). In this study, our data showed that GCE prevented calcium oxalate kidney stone formation in *Drosophila* renal tubules. We also showed that treatment with GCE completely removed pre-formed calcium oxalate kidney stones from adult *Drosophila* renal tubules in a concentration-dependent manner. Surprisingly, HCA added in fly food had no effect on the formation and removal of calcium oxalate renal stones in *Drosophila* Malpighian tubules even at a very high concentration. We speculated that HCA added in fly food could not be absorbed and transported to Malpighian tubules because of the absence of key components in GCE which were essential for the delivery of HCA to Malpighian tubules. Our study also showed that GCE was effective in both genetic and non-genetic *Drosophila* kidney stone models, suggesting that GCE could be a novel medication for the treatment of nephrolithiasis. Nevertheless, our study showed that the *Drosophila* Malpighian tubule nephrolithiasis model could be used to efficiently screen thousands of traditional Chinese herbal medicine (TCM) to seek the golden treasure of TCM for nephrolithiasis treatment.

Our study also showed that hydroxycitrate could directly dissolve calcium oxalate stones in renal tubules *ex vivo* in a concentration-dependent manner. The efficiency of HCA dissolving calcium oxalate stones in renal tubule is very high. Calcium oxalate stones in dissected renal tubules were completely dissolved in 0.5% and 1% HCA within 120 minutes. On the other hand, we thought that the HCA concentration in 2% GCE would be high enough for the dissolution of calcium oxalate stones because GCE contains 60% HCA. However, 2% GCE surprisingly had no effect on the dissolution of calcium oxalate stones in Malpighian tubules *ex vivo.* We reasoned that this was because 31.3% of GCE is composed of fiber, which cannot be dissolved in water, and as a result, HCA could not be released from GCE water solution. We also tried GCE solution in ethanol, but it also did not dissolve calcium oxalate stones *ex vivo*. We did not observe any impact of GCE on the removal of calcium oxalate stones *ex vivo*, suggesting that the release of HCA from GCE is essential for its function to remove the calcium oxalate stones *in vivo*. GCE added in fly food could be absorbed in midgut and released into hemolymph as free HCA form, and then secreted into Malpighian tubule lumen where HCA could dissolve calcium oxalate renal stones．Our results strongly support the idea that *Garcinia cambogia* extract removes calcium oxalate renal stones via the direct dissolution of stones by HCA in the renal tubules.

Currently potassium citrate is often prescribed to patients with calcium oxalate kidney stone diseases. However, its clinical impact on the removal of kidney stones is still controversial. In this study, we showed that citric acid efficiently dissolved calcium oxalate stone in dissected renal tubules *ex vivo*, however, the effect of citric acid on calcium oxalate kidney stone is minimal compared to GCE *in vivo*. To improve the efficiency of citric acid in clinic, we need to find better ways to deliver potassium citrate to patients.

In summary, this work is the first to clearly demonstrate that clinical-grade *Garcinia cambogia* extract removes calcium oxalate stones in *Drosophila* Malpighian tubule *in vivo* and hydroxycitrate can directly dissolve calcium oxalate stones in renal tubules *ex vivo*. Our study discovered a novel therapeutic strategy for the clinical treatment of nephrolithiasis, and clinical-grade *Garcinia cambogia* extract could be used to treat patients with nephrolithiasis in the near future.

### CONCLUSIONS

*Garcinia cambogia* extract removes calcium oxalate stones in *Drosophila* Malpighian tubule, and hydroxycitrate-a major component of GCE, directly dissolve calcium oxalate stones in renal tubules *ex vivo*. *Garcinia cambogia* extract has the potential to be used to treat patients with nephrolithiasis.

## METHODS

### Fly Strains

Flies were reared on standard food at 25°C. All UAS-Gal4 crosses were performed at 25°C. Uro-Gal4 (Bl-44416) and UAS-nGFP (Bl-4775) were obtained from the Bloomington *Drosophila* stock center. UAS-RNAi transgenic fly lines targeting vha100-2 (TH04790.N, Bl-64859) and vha55 (THU4117 and v-46554), referred in the main text and figures as vha100-2 IR and vha55 IR, were obtained from the Bloomington *Drosophila* stock center, Vienna stock center and Tsinghua Fly center. Uro-Gal4 and UAS-nGFP were recombined together to label principal cells at all developmental stages.

### Chemicals

Potassium hydroxycitrate tribasic monohydrate (59847) and Potassium citrate tribasic monohydrate (c8385) were purchased from Sigma. Swanson^®^ Super Citrimax Clinical Strength *Garcinia Cambogia* was purchased online from Amazon. Other chemicals were purchased from Sigma unless otherwise indicated.

### RNAi knockdown of nephrolithiasis-related genes in principal cells

This method was adopted from our recent study described previously(Zhang et al., 2013). Briefly, the UAS/Gal4 system allows for the over-expression or knockdown of “gene-of-interest” in a cell-specific manner in *Drosophila*. A fly line that possesses a “GAL4-driver” can be crossed to a second transgenic fly line containing a construct of interest gene X placed downstream of a UAS promoter sequence. This allows the downstream transgene to be expressed specifically in these cells where GAL4 is expressed. To specifically knock down nephrolithiasis-related genes in Malpighian tubule principal cells, we used the Uro-GAL4 driver, which drives gene expression specifically in the tubular principal cells, and crossed it to the UAS-RNAi lines containing a dsRNA hairpin directed against gene X.

### RNAi -based functional analysis of malpighian tubule genes

10 virgins of Uro-Gal4/UAS-nGFP flies were crossed with 5 males of UAS-RNAi transgenic line in vials at 25°C. Freshly hatched flies were transferred to fly food with 0.3% NaOx at 25°C for 1 week. Malpighian tubules were dissected in PBS under dissection microscope and then subjected to the examination of renal stone in malpighian tubules under polarized white light with an Olympus BX63 optical microscope. We randomly selected three images for quantification. Renal stone formation was measured in the whole field of view (700umx100um, 20x magnification) using cellSens software. The results were expressed as mean ± SD (n=10). All statistical analyses were performed using GraphPad Prism5 software. Statistical significance was defined as P<0.05.

### Calcium oxalate stone prevention analysis *in vivo*

Briefly, wild type or mutant flies were reared on regular fly food containing 0.1% NaOx and different concentration of HCA or GCE at 25°C for one week. Malpighian tubules of adult female flies were dissected in PBS under dissection microscope and then subjected to the examination of renal stone in malpighian tubules under polarized white light with an Olympus BX63 optical microscope. We randomly selected three images for quantification. Renal stone formation was measured in the whole field of view (700umx100um, 20x magnification) using cellSens software. The results were expressed as mean ± SD (n=10). All statistical analyses were performed using GraphPad Prism5 software. Statistical significance was defined as P<0.05.

### Calcium oxalate stone removal assay *in vivo*

First, wild type or mutant flies were reared on regular fly food containing 0.3% NaOx at 25°C for one week and renal tubule stone formation was evaluated briefly. Then, flies were transferred to fly food containing 0.1% NaOx and different concentration of HCA or GCE at 25°C for 1 week. Malpighian tubules of adult female flies were dissected in PBS under dissection microscope and then subjected to the examination of renal stone in malpighian tubules under polarized white light with an Olympus BX63 optical microscope. We randomly selected three images for quantification. Renal stone formation was measured in the whole field of view (700umx100um, 20x magnification) using cellSens software. The results were expressed as mean ± SD (n=10). All statistical analyses were performed using GraphPad Prism5 software. Statistical significance was defined as P<0.05.

### Ex vivo Calcium oxalate stone dissolution analysis

Briefly, wild type w^1118^ flies were reared on regular fly food containing 0.3% NaOx for one week. Intact Malpighian tubules were dissected in PBS under dissection microscope and transferred onto a slide. 100ul CA, HCA or GCE solution was added to completely cover the MT tissue, and then the Malpighian tubules were subjected to live-imaging under polarized white light with an Olympus BX63 optical microscope without a coverslip. Images were taken every 20min and the total area of renal stones was measured using cellSens software. Renal stone dissolution rate was calculated by dividing the remaining stone area by the total stone area at the beginning. The results were expressed as mean ± SD (n=5). All statistical analyses were performed using GraphPad Prism5 software.

## LIST OF ABBREVIATIONS

GCE: *Garcinia cambogia* extract
HCA: hydroxycitric acid
CaOx: calcium oxalate
NaOx: Sodium oxalate
MsT: Malpighian tubule

## Authors’ contributions

F.J.Z. and F.F.H. designed the study; Q.X.F., X.M.F., X.Z.H, S.Q.G and J.W.T carried out the experiments; Q.X.F., F.F.H and F.J.Z. analyzed the data; Q.X.F. and F.J.Z. made the figures; F.F.H and F.J.Z. drafted and revised the paper; all authors approved the final version of the manuscript.

## Acknowledgements

We thank the Bloomington fly stock center and the Tsinghua fly stock center for *Drosophila* stocks. We want to thank Dr. Haiyan Fu for buying and bringing back GCE reagents from US.

---

**FigureS1.**
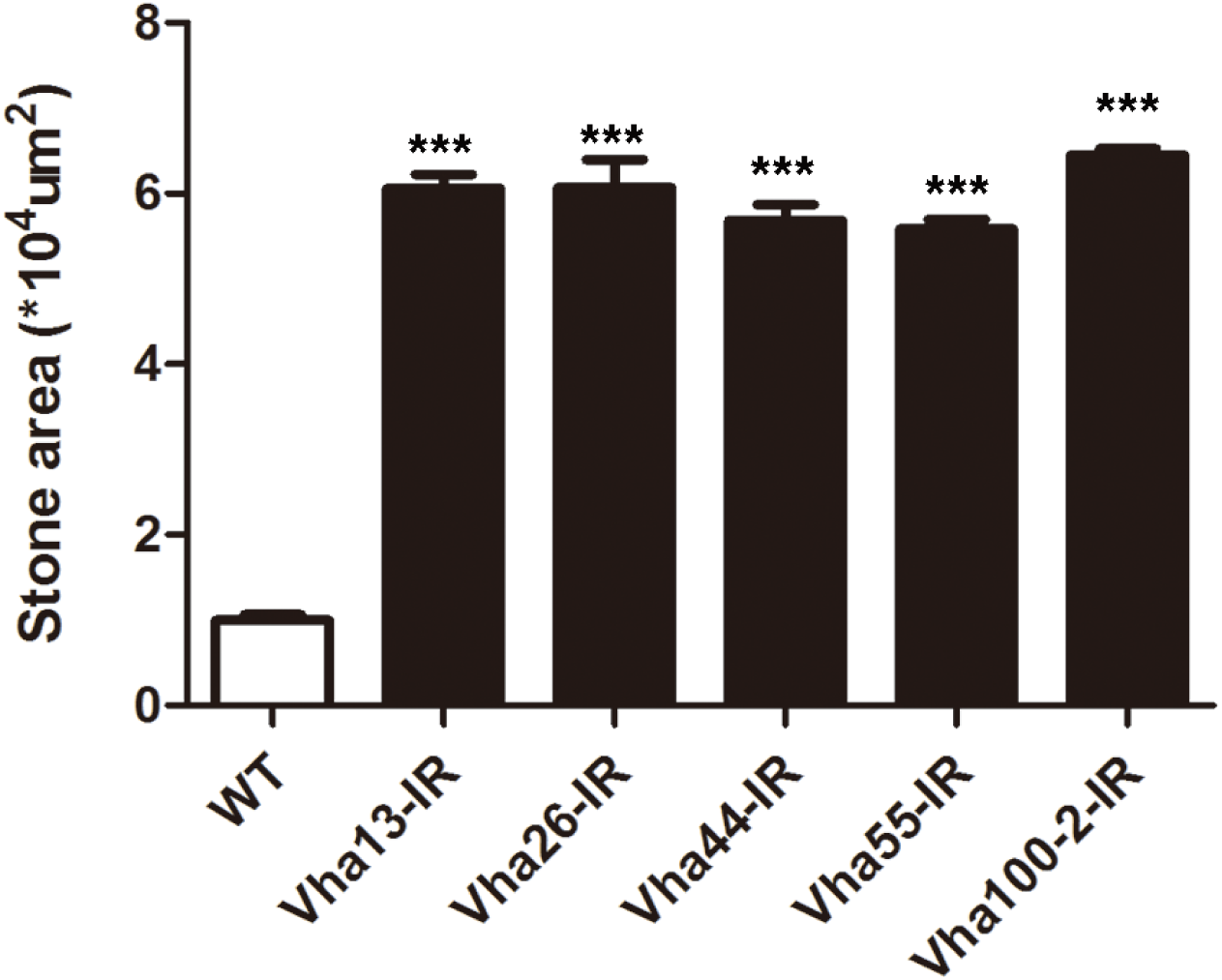
The effects of fly orthologs of 6 mammalian v-ATPase genes on CaOx stone formation in Malpighian tubules. RNAi Knock-down of each gene of v-ATPase complex led to increased formation of calcium oxalate stone in *Drosophila* Malpighian tubule. The results are expressed as mean ± SD. The unpaired T-test was performed with two-tailed p-values and 95% confidence intervals. Statistical significance was defined as P<0.05. (*p<0.05, **p<0.005, ***p<0.0005).

